# Obliquity Feature Extraction for Fossil Data Analysis: The Stickleback Fish Case

**DOI:** 10.64898/2026.02.01.703096

**Authors:** Rolf Ergon

## Abstract

A moving average smoothing method for extraction of cycles in time series data is described, with focus on obliquity cycles and fossil data. The proposed method is intended for cases where the environmental driver of phenotypic evolution can be shown to include obliquity cycles, either by power spectrum analysis or simply by inspection of raw or smoothed time series. The method gives improved mean trait predictions and better understanding when applied on stickleback fish fossil data from around 10 million years ago. The possibility to extract obliquity cycles will depend on the dynamics of the time series, and the method is thus not universally applicable. It may, however, be possible to adapt the size of the moving window to problems under study, or possibly to obtain improved predictions by inclusion of a sinusoidal component in the mean trait prediction modeling.

## 1 Introduction

Obliquity refers to the angle of tilt of a planet’s rotational axis relative to its orbital plane, and on Earth this angle varies a few degrees with cycles of around 41,000 years (41 kyr) (Wikipedia contributors, 2025). Information on obliquity cycles is utilized in climate and oceanographic research, and the more precise value of the period varies in the literature. Tian et al. (2013) refer to both 40 and 41 kyr around 14 million years ago (Ma), Zeebe and Lourens (2022) use 40.5 kyr around 40 Ma, and Sullivan et al. (2025) use 40.3 kyr around 18 Ma. Based on power spectrum analysis in Section 3 below, I here use 40.6 kyr.

Until recently obliquity cycles were apparently not referred to in fossil data studies. In Ergon (2025b), however, I showed that the phenotypic evolution of a trait in a stickleback fish lineage around 10 Ma was partly driven by obliquity cycles. This is an example of adaptive peak tracking as described in Ergon (2025a), with oxygen isotope *∂*^18^*O*(*t*) values used as environmental driver. Since these cycles affect marine life, they have left traces in the *∂*^18^*O*(*t*) values over time, as found in deep-sea drilling projects (Westerhold et al., 2020). The underlying connection between deep-sea drilling data and the evolution of stickleback fish traits as described in Bell et al. (1985) is that the obliquity cycles also affected the evolution of these fishes.

The extraction of obliquity information from *∂*^18^*O*(*t*) data is described in Appendix B in Ergon (2025b), and the purpose of the present article is to give a more thorough explanation. In Appendix B in Ergon (2025b), the feature extraction method was also illustrated by power spectrum analysis, which unfortunately had some serious errors. This is corrected here. As will be shown, the power spectrum of *∂*^18^*O*(*t*) time series gives in fact the fundamental explanation of the improved stickleback fish results in Ergon (2025b), as compared with earlier results.

The theoretical background for the feature extraction method is given in Section 2, while the detailed explanation is given in Section 3, using the stickleback fish case as example. A summary with discussion follows in Section 4. An example of non-linear estimation of the adaptive peak function is given in Appendix A, and information scaling aspects are visualized in Appendix B. MATLAB code and oxygen isotope data are archived at bioRxiv.

## 2 Theory and Methods

### 2.1 Extraction of obliquity information

For extraction of obliquity information, the *∂*^18^*O*(*t*) time series as found in, e.g., Westerhold et al. (2020) can be modeled as

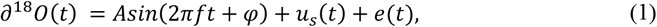

where *f* = 24.7 periods per million year (Myr). Here *u*_*s*_(*t*) includes structural information that is important for prediction of the mean traits under study, while *e*(*t*) is unimportant high frequence noise. Also assume a sampling frequency *f*_*s*_ ≫ *f*, and centered moving average smoothing with rectangular window and sample size *n*, where *n* is large enough to make the smoothed high frequency noise *e*_*smooth*_(*t*) ≈ 0.

For an integer number of cycles in the moving time window, the average of the sinusoidal component in Eq. (1) is zero. For a non-integer number of periods in the window we thus find

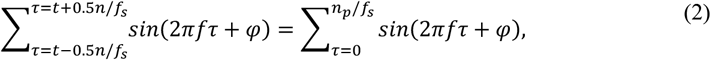

where 0 < *n*_*p*_ < *f*_*s*_/*f*, with *f*_*s*_/*f* being the number of samples per period. From Eq. (1) we thus find

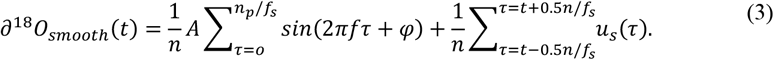

For a non-integer number of periods in the moving time window, the sinusoidal component will thus appear in *∂*^18^*O*_*smooth*_(*t*) with the frequency *f*, but with an amplitude depending on *n* and *n*_*p*_ according to Eq. (3), while the structured information will appear as *u*_*s,smooth*_(*t*). The sampling window must thus be wide enough to filter out the noise component *e*(*t*), and narrow enough to retain some scaled information on both the sinusoidal component and essential features in the structured information in *u*_*s*_(*t*). The first term in Eq. (3) will in theory have local maxima for time window sizes *T*_*i*_ = (*i* + 0.5) · 40.6 kyr, where *i* is an integer number, but with decreasing values for increasing window sizes, and the value of *T*_*i*_ for best mean trait predictions can be found from weighted mean squared errors, or alternatively from Akaike information criterion values. This value of *T*_*i*_ is not necessarily the optimal window size because also *u*_*s,smooth*_(*t*) is an important part of *∂*^18^*O*_*smooth*_(*t*), and it may thus be possible to find an even better window size by tuning for optimal predictions.

### 2.2 Other methods

Weighted least squares (WLS) estimation and Akaike information criterion (*AIC*) computations will be used as described in Ergon (2025b). The *AIC* values depend on the number of estimated parameters, including those found by manual searches, and in the stickleback fish case they must also be compensated for short data, such that *AIC*_*c*_ must be used.

## 3 The stickleback fish case used as example

Data for the stickleback fish case in Bell et al. (1985) are summarized in Ergon (2025b), while *∂*^18^*O*(*t*) from 9,000 to 11,200 Ma are found in Westerhold et al. (2020), Table S34, column marked `ISOBENbinned_d18Ointerp`. The sampling frequency in these data is *f*_*s*_ = 500 samples per Myr, and we thus have *f*_*s*_⁄*f* ≈ 20 samples per obliquity cycle.

The zero time point is not given in Bell et al. (1985), but since *t* = 0 in the stickleback fish data is approximately 10 Ma (personal information) it was possible to align the data for optimal mean trait predictions, and the best tracking result was obtained when *t* = 0 in the stickleback fish data corresponds to *t*_0_ = −9.994 Myr with present time as zero point (Ergon, 2025b).

The true *∂*^18^*O*(*t*) time series is shown in Fig. 1, upper panel, together with a moving mean time series *∂*^18^*O*_*mean*_(*t*) obtained from a centered and rectangular time window of 81 kyr. With this window size of two obliquity cycles, the cycles will according to Eq. (3) not be seen in *∂*^18^*O*_*mean*_(*t*). Fig. 1, lower panel, shows simulated data

**Figure 1.**
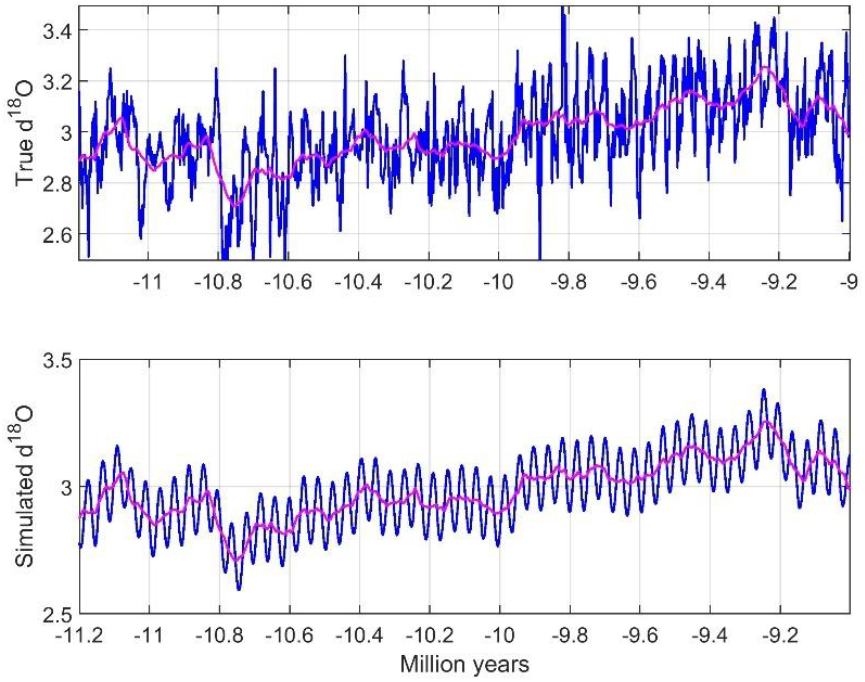
True *∂*^18^*O*(*t*) and moving mean *∂*^18^*O*_*mean*_(*t*) time series (upper panel, blue and red lines), and simulated time series according to Eq. (4) (lower panel).

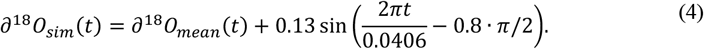

with amplitude *A* = 0.13 and phase shift *φ* = −0.8 · *π*/2 chosen such that mean trait predictions based on *∂*^18^*O*_*sim*_(*t*), as explained below, had minimum WMSE. As shown in Fig. 3, the simulated cycles were then also aligned with the true cycles.

It is far from easy to see in Fig. 1 that the true *∂*^18^*O*(*t*) signal includes obliquity cycles. This is, however, found by means of power spectrum analysis using the function periodogram in MATLAB, and for that purpose variations around the moving mean values as shown by red lines in Fig. 1 were used. Because there are 1101 samples in the raw data, Fourier analysis over 1024 frequency values were then possible. Fig. 2 shows that the true *∂*^18^*O*(*t*) signal after removal of *∂*^18^*O*_*mean*_(*t*) has a significant spectral density peak with a frequency of 24.7 periods per Myr, corresponding to the obliquity cycles. As the simulated signal after removal of *∂*^18^*O*_*mean*_(*t*) is a pure sinus signal, the power spectrum is a narrow single peak at 24.7 periods per Myr.

**Figure 2.**
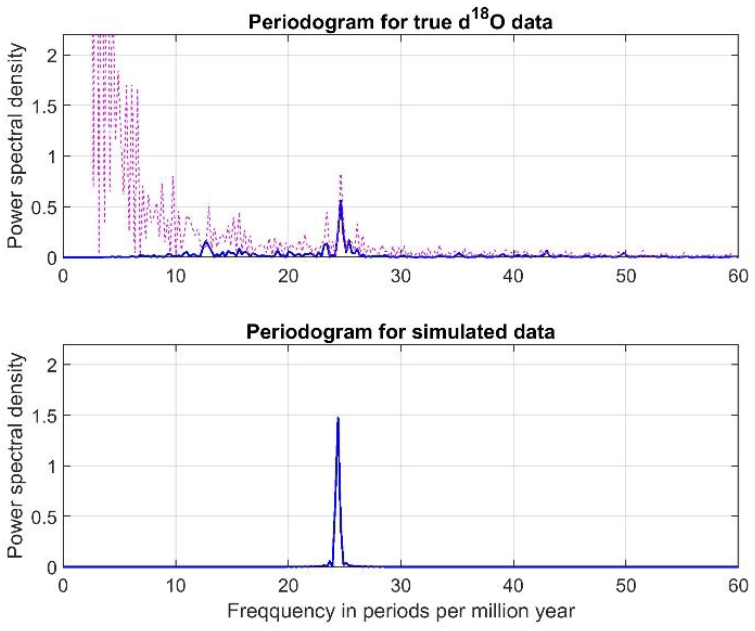
Periodograms for *∂*^18^*O*(*t*) and *∂*^18^*O*_*sim*_(*t*) after subtraction of moving mean values (blue lines). Dashed red line shows the periodogram for the true time series, without subtraction of moving mean values.

**Figure 3.**
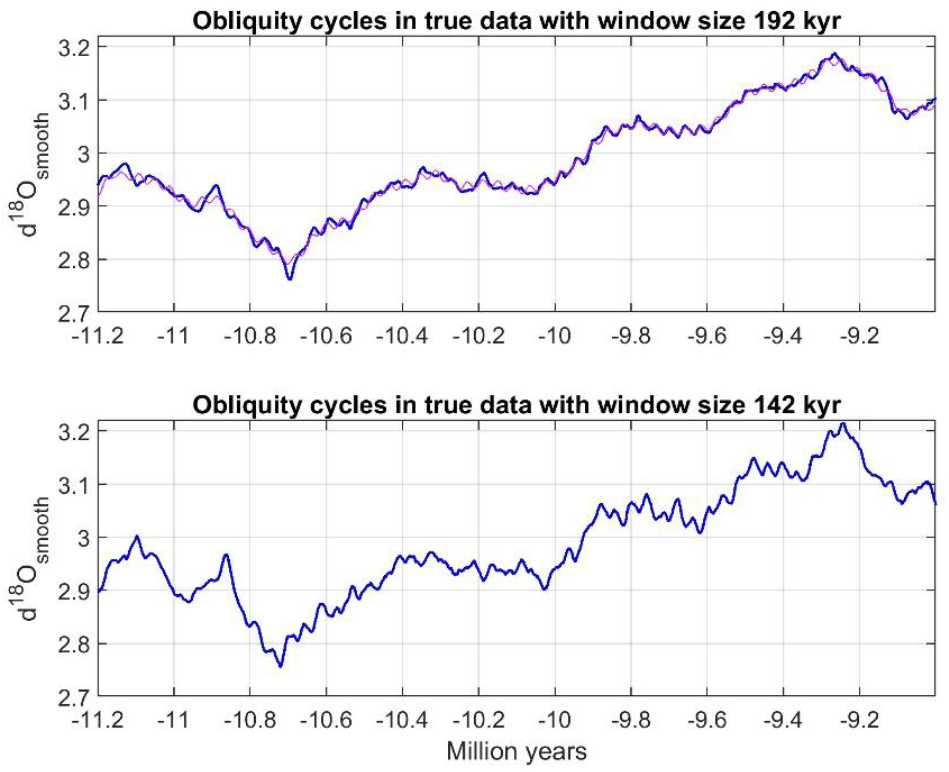
Extracted obliquity cycles using moving average smoothing of the true data with two different window sizes (blue lines). Upper panel also shows time series with simulated cycles after smoothing (red line).

Among moving window sizes of 1.5, 2,5, 3.5, 4.5 and 5.5 obliquity cycles, the best stickleback fish mean trait predictions are found by use of 4.5 cycles, which since *∂*^18^*O*(*t*) has 500 samples per Myr corresponds to window size 183 kyr. With this as starting point, the optimal window size giving the lowest WMSE value was found to be 192 kyr. The smoothed true signal *∂*^18^*O*_*smooth*_(*t*) using this window size is shown in Fig. 3, upper panel, where also the smoothed version *∂*^18^*O*_*sim,smooth*_(*t*) of *∂*^18^*O*_*sim*_(*t*) is shown. Note that the obliquity cycles are clearly visible only in some time periods, and that the stickleback fish data falls within such a period around 10 Ma. As shown in Fig. 3, lower panel, a window size of 3.5 · 40.6 = 142 kyr could possibly be a good choice for other cases.

Results for the stickleback fish adaptive peak estimation and mean trait predictions, as discussed in more detail in Ergon (2025a,b), are shown in Figures 4 and 5. The adaptive peak function in Fig. 4 was here found to be linear, but as shown in Appendix A it may also be non-linear. The predictions based on true data with the optimal moving window size *T* = 192 kyr resulted in *WMSE* = 2.11 · 10^−4^ (*k* = 5, *AIC*_*c*_ = −131). The data with simulated cycles according to Eq. (4) with use of the same window size resulted in *WMSE* = 2.31 · 10^−4^ (*k* = 7, *AIC*_*c*_ = −122). For comparison, the WLS predictions as shown by straight line in Fig. 5 has *WMSE* = 4.01 · 10^−4^ (*k* = 3, *AIC*_*c*_ = −121). These results for the various models are summarized in Table 1, including Akaike weights, which can be directly interpreted as conditional probabilities for the models (Wagenmakers and Farrell, 2004). Here, results for the sinusoidal model in Ergon (2025a), without use of *∂*^18^*O*(*t*) data, are included. Note that the model with simulated cycles has much lower WMSE than the WLS model, but that owing to the number of estimated parameters it has almost the same *AIC*_*c*_ value. Also note that the sinusoidal model without us of *∂*^18^*O*(*t*) data has the lowest WMSE and *AIC*_*c*_ values, but that the true *∂*^18^*O*(*t*) data model is nearly as good.

**Table 1.**
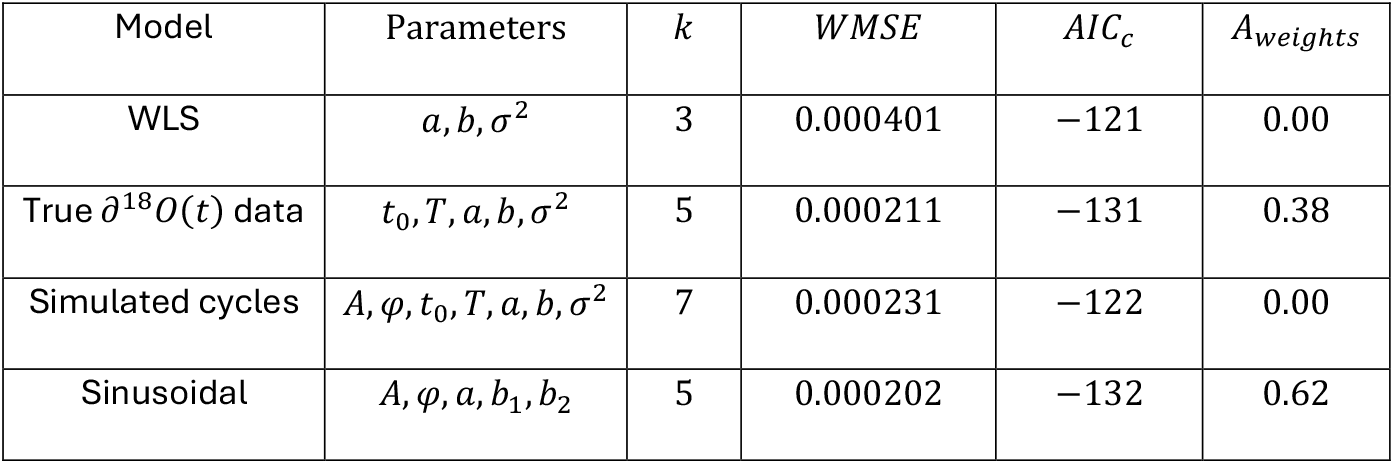
Model comparisons.

**Figure 4.**
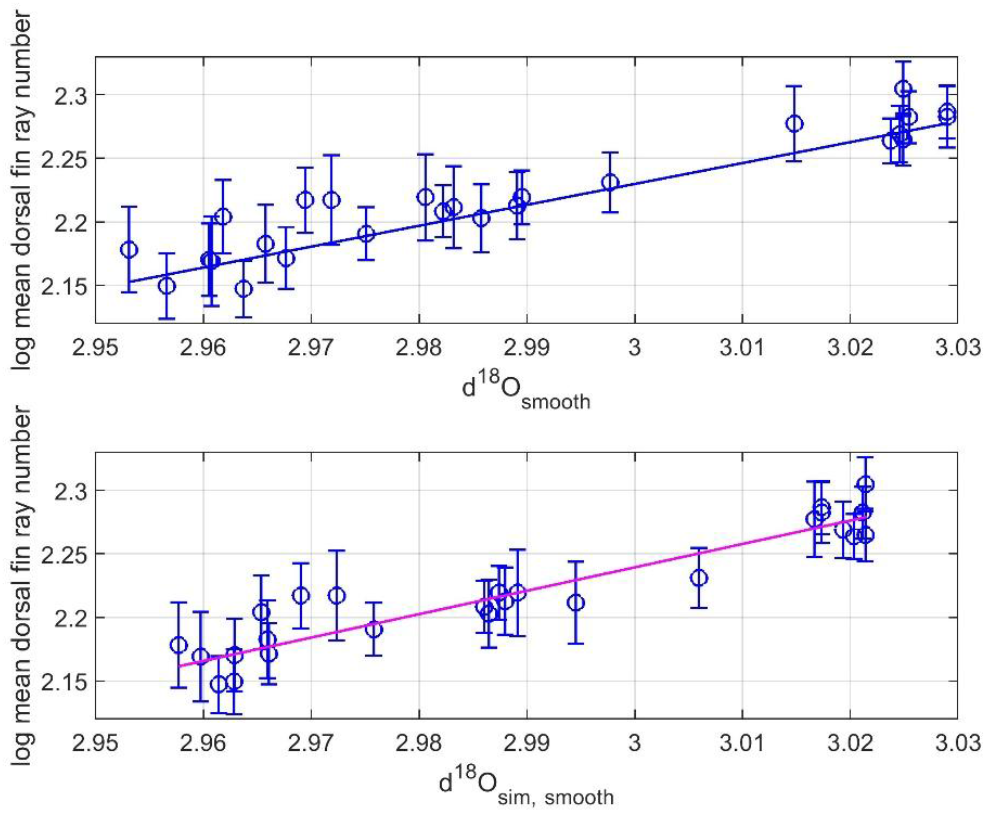
Mean trait values 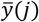 with standard error bars versus environmental driver values *∂*^18^*O*_*smooth*_(*j*) (upper panel) and *∂*^18^*O*_*sim,smooth*_(*j*) (lower panel), for *j* = 1, 2, …, 26, and estimated linear adaptive peak functions (blue and red lines).

**Figure 5.**
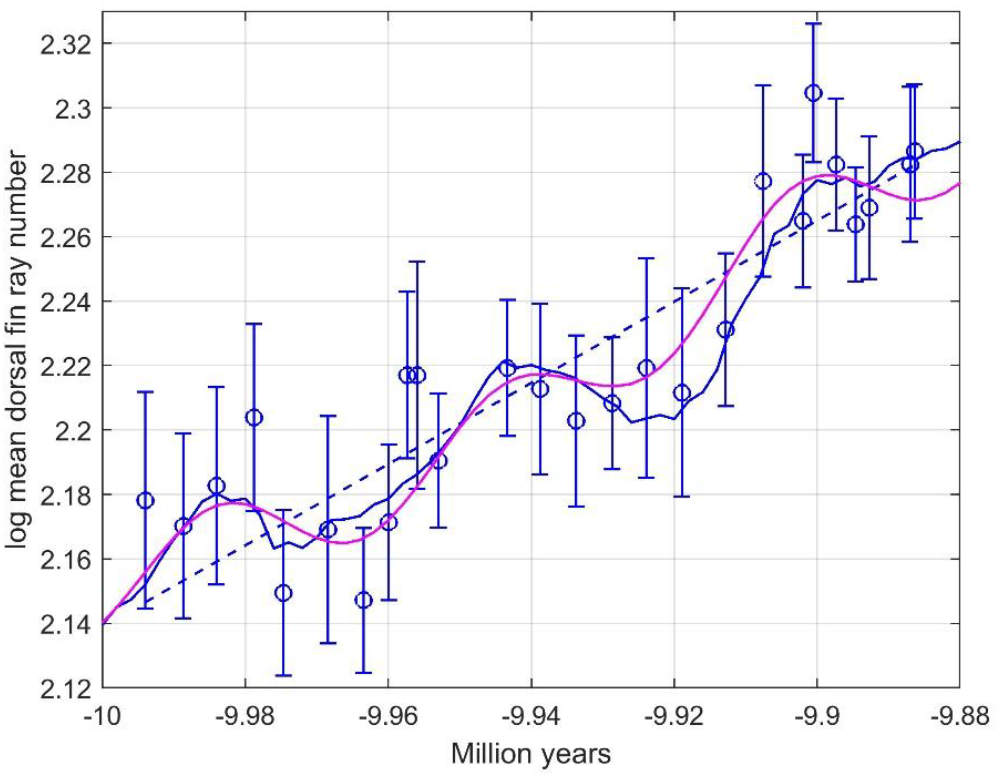
Predictions of stickleback fish mean trait by use of true data (blue line), and by use of data with simulated obliquity cycles (red line). The moving window size was 192 kyr. Dashed line shows WLS predictions.

## 4 Summary and discussion

The feature extraction method used in Ergon (2025b) is described in some more detail, and also clarified by use of simulations. As apparent from Fig. 1, it is far from obvious that the *∂*^18^*O*(*t*) time series obtained from deep-sea drilling samples contains obliquity cycles. This is, however, shown by the spectral analysis results in Fig. 2, with a peak for the true signal that corresponds to a period of 40.6 kyr. The power spectrum also has a minor and unknown frequency component corresponding to a period of 77 kyr.

The obliquity cycles in *∂*^18^*O*_*smooth*_(*t*) used for predictions are seen in Fig. 3, upper panel. As follows from Eq. (3), both the sinusoidal component and the structured information *u*_*s*_(*t*) are scaled in the same way by the number of samples in the moving window, and this scaling is accounted for in the adapted peak function as found in Fig. 4. The effect of this scaling is visualized in Appendix B.

As shown in Table 1 and Fig. 5, the proposed feature extraction method is quite useful in the stickleback fish case, as also discussed in Ergon (2025b). The *∂*^18^*O*(*t*) signal obtained through deep-sea drilling can thus be used as proxy for the temperature at the field site in what is now Nevada, scaled in an appropriate way and presumably during a critical period of the year.

Centered moving average smoothing may thus extract obliquity cycles for the purpose of mean trait predictions, but as shown in Fig. 3 this is not a universally reliable method. As also shown in Fig. 3, however, the size of the moving average window could possibly be adapted to prediction problems under study.

Realizing that the mean traits may be influenced by obliquity cycles, as indicated by a power spectrum as in Fig. 2, simulated cycles may be included in the modeling, as discussed in Section 3 and with prediction results as shown in Table 1 and Fig. 5. Although the partly simulated model gives almost as good predictions as the true data model, the *AIC*_*c*_ value is poor owing to the increased number of estimated parameters. In that respect a sinusoidal model based on assumed obliquity cycles, and without use of *∂*^18^*O*(*t*) data, is better (Ergon, 2025b). As seen in Table 2 this is in fact the best model with the most negative *AIC*_*c*_ value and Akaike weight 0.62, but this result depends on the number of parameters needed for modeling of the structured information *u*_*s*_(*t*) in Eq. (1). Also note that the model based on *∂*^18^*O*(*t*) data with Akaike weight 0.38 is almost as good.

The main conclusion from the findings in Ergon (2025b) and in this article is that sinusoidal models as in Ergon (2025a), without use of *∂*^18^*O*(*t*) data, and models based on true and smoothed *∂*^18^*O*(*t*) data may be equally good, and that the choice must be made in each specific case.

It remains to be seen if the methods discussed in this article, and successfully applied on the stickleback fish case, can give improved predictions in other cases where the presence of obliquity cycles are found by use of power spectrum analysis or simply by inspection of raw or smoothed time series.

## Supporting information

MATAB Code

## Acknowledgements

I thank Mike Bell for information regarding the time span of the stickleback fish data and for the observation of possible obliquity cycles. I also thank University of South-Eastern Norway for support.

## Data Availability Statement

MATLAB code and oxygen isotope data are archived at bioRxiv, https://doi.org/10.64898/2026.02.01.703096

## A. Non-linear adaptive peak function

For a window size of 183 kyr, i.e., of 4.5 obliquity cycles, the adaptive peak function becomes non-linear, and as shown in Fig. A, upper panel, it may then be modeled as a second order function. Such a non-linear adaptive peak function raises an interesting question. Did obliquity cycles at the field site affect the adaptive peak through a non-linear function, or does the use of *∂*^18^*O*(*t*) values obtained through deep-sea drilling represent a temperature measurement error with non-linearity as consequence? Since the adaptive peak function for the simulated data model is almost linear, the non-linearity appears to be caused by unexplained variations in the true time series, i.e., a form of measurement error. This argument is strengthened by the result in Fig. 4, where both adaptive functions appear to be linear.

**Figure A.**
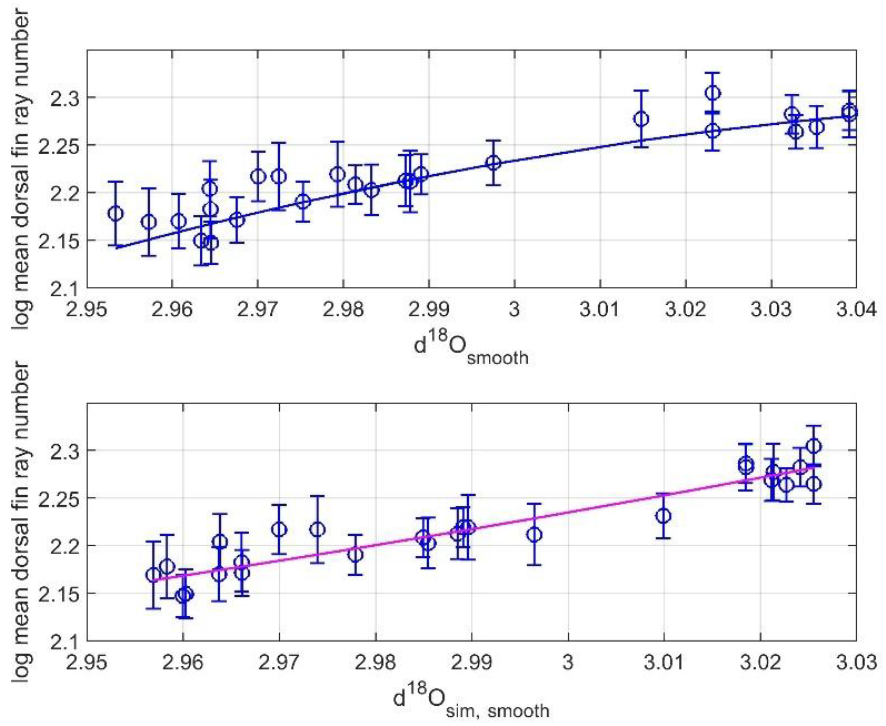
Mean trait values 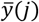 with standard error bars versus environmental driver values *∂*^18^*O*_*smooth*_(*j*) (upper panel) and *∂*^18^*O*_*sim,smooth*_(*j*) (lower panel), for *j* = 1, 2, …, 26, and estimated non-linear adaptive peak functions (blue and red lines). The time window size was 183 kyr.

## B. Information scaling

As follows from Eq. (3), both the sinusoidal component and the structured information *u*_*s*_(*t*) are scaled in the same way by the number of samples in the moving window. This is illustrated in Fig. B., showing *∂*^18^*O*_*smooth*_(*t*) for time window sizes 192 kyr and 2.5 · 40.6 = 102 kyr. The prediction results for these window sizes are *WMSE* = 2.11 · 10^−4^ (*k* = 5, *AIC*_*c*_ = −131) and *WMSE* = 3.73 · 10^−4^ (*k* = 5, *AIC*_*c*_ = −117), respectively. For comparison, the WLS predictions as shown by dashed straight line in Fig. 5 has *WMSE* = 4.01 · 10^−4^ (*k* = 3, *AIC*_*c*_ = −121).

**Figure B.**
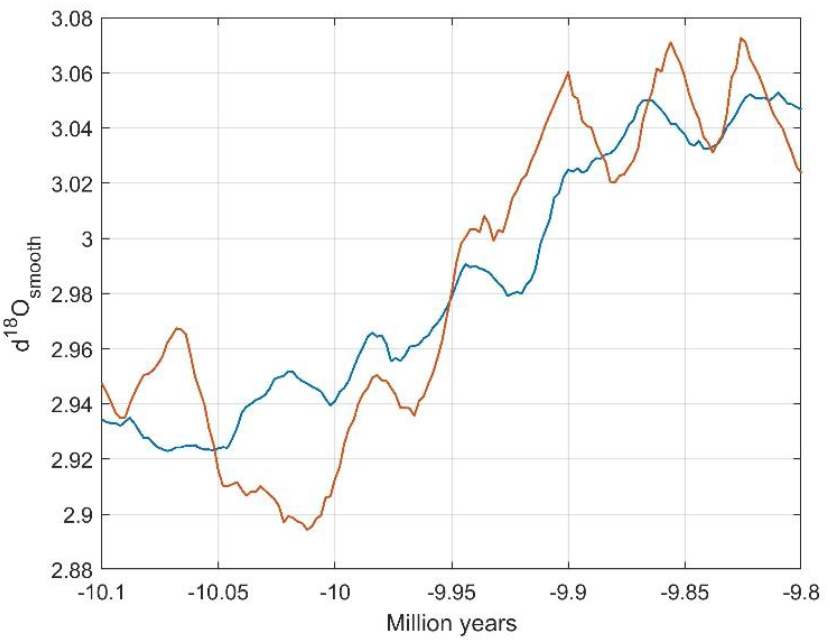
Segments of *∂*^18^*O*_*smooth*_(*t*) for window sizes 192 and 102 kyr, shown by blue and red lines, respectively.

## References

Bell, M.A., Baumgartner, J.V., and Olson, E.C. Patterns of temporal change in single morphological characters of a miocene stickleback fish. Paleobiology, 1985. 11(3):258–271. doi:10.1017/S009483730000590X.

Ergon, R. Adaptive Peak Tracking as Explanation of Sparse Fossil Data Across Fluctuating Ancient Environments. Ecology and Evolution, 2025a. 10.1002/ece3.71705.

Ergon, R. Examples of Adaptive Peak Tracking as Found in the Fossil Record, Including Obliquity Cycles Tracking. Modeling, Identification and Control, 2025b. doi:10.4173/mic.2025.4.2.

Sullivan et al. Obliquity disruption and Antarctic ice sheet dynamics over a 2.4-Myr astronomical grand cycle. Sci. Adv., 2025. doi:10.1126/sciadv.adl1996.

Tian, J., Yang, M., Lyle, M.W., Wilkens, R. and Shackford, J.K. Obliquity and long eccentricity pacing of the Middle Miocene climate transition. Geochemistry, Geophysics, Geosystems, 2013. doi:10.1002/ggge.20108.

Wagenmakers, E.-J. and Farrell, S. AIC model selection using Akaike weights. Psychonomic Bulletin & Review, 2004. 11(1):192–196. doi:10.3758/BF03206482.

Westerhold, T. et al. An astronomically dated record of earth’s climate and its predictability over the last 66 million years. Science, 2020. 369(6509):1383–1387. doi:10.1126/science.aba6853.

Wikipedia contributors. Milankovitch cycles. https://en.wikipedia.org/wiki/Milankovitch_cycles, 2025. [Online; accessed January-2026].

Zeebe, R.E. and Lourens, L.J. A Deep-Time Dating Tool for Paleo-Applications Utilizing Obliquity and Precession Cycles: The Role of Dynamical Ellipticity and Tidal Dissipation. Paleoceanography and Paleoclimatology, 2022. doi:10.1029/2021PA004349.

