## Supplementary material for "Obliquity Feature Extraction for Fossil Data Analysis: The Stickleback Fish Case": MATAB Code

### MATLAB code for

### The Stickleback Fish Case

Rolf Ergon

University of South-Eastern Norway

February 9, 2026

```
clear

%% True data

load Data_9000_to_11200.mat

Data=[Age d18O];

Age=flipud(Age);
d18O=flipud(d18O);
tdata=-Age';            % Step size 0.002 My
utrue=d18O';
umean=movmean(utrue,41);

%% Simulated data
usin=sin(2*pi*tdata/0.041-0.8*pi/2);
usim=umean+0.13*usin;
```

```
figure(1)
subplot(2,1,1)
plot(tdata,utrue,'b','LineWidth',1.0), hold on
plot(tdata,umean,'m','LineWidth',1.0), hold off, grid
axis([-11.2 -9 2.5 3.5])
ylabel('True d^1^8O')
subplot(2,1,2)
plot(tdata,usim,'b','LineWidth',1.0), hold on
plot(tdata,umean,'m','LineWidth',1.0), hold off, grid
axis([-11.2 -9 2.5 3.5])
ylabel('Simulated d^1^8O')
xlabel('Million years')

%% Spectral analysis
ures_true=utrue-umean;
ures_true_raw=utrue;

ures_sim=usim-umean;
ptrue=periodogram(ures_true);
ptrue_raw=periodogram(ures_true_raw);
psim=periodogram(ures_sim);
fplot=0:1024;
fplot=fplot/4.096;

figure(2)
subplot(2,1,1)
plot(fplot,ptrue','b','LineWidth',1.0), hold on
plot(fplot,ptrue_raw','--m'), hold off, grid
title('Periodogram for true d^1^8O data')
ylabel('Power spectral density')
axis([0 60 0 2.2])
subplot(2,1,2)
plot(fplot,psim','b','LineWidth',1.0), grid
```

```
title('Periodogram for simulated data')
xlabel('Freqquency in periods per million year')
ylabel('Power spectral density')
axis([0 60 0 2.2])

window=96;

% window=3.5*40.6/2;      % For. Fig. 3, lower panel

% window=2.5*40.6/2;      % For Fig. B

utrue_smooth=movmean(utrue,window);
usim_smooth=movmean(usim,window);

figure(3)
subplot(2,1,1)
plot(tdata,utrue_smooth,'b','LineWidth',1.0), hold on
plot(tdata,usim_smooth,'m'), hold off, grid
title('Obliquity cycles in true data with window size 192 kyr')
ylabel('d^1^8O_s_m_o_o_t_h')
axis([-11.2 -9 2.7 3.22])

window2=3.5*40.6/2;
utrue_smooth2=movmean(utrue,window2);
subplot(2,1,2)
plot(tdata,utrue_smooth2,'b','LineWidth',1.0), grid
title('Obliquity cycles in true data with window size 142 kyr')
xlabel('Million years')
ylabel('d^1^8O_s_m_o_o_t_h')
axis([-11.2 -9 2.7 3.22])

%% Stickleback sample data
N=26;
k=5;
```

```
t0=-10+0.006;
tlabel1=[0 5317 9950 15182 19318 25551 30541 34030 36631 38052 41008 50629 55246];
tlabel2=[60289 65312 70119 75075 81150 86454 92006 93445 96659 99377 101306 106969 107667];
tlabel=[tlabel1 tlabel2]/1000000;
tlabel=t0+tlabel;
n=[30 33 30 36 55 40 75 52 22 28 54 45 35 37 41 25 30 58 32 84 82 89 95 57 45 64];
y1=0.01*[883 876 887 906 858 875 856 877 918 918 894 920 914 ];
y2=0.01*[905 910 920 913 931 975 963 1002 980 962 967 980 984];
y=[y1 y2];
sd1=0.001*[747 663 681 715 762 899 775 703 501 772 627 588 648  ];
sd2=0.001*[664 539 707 730 754 718 803 846 828 732 715 694 718];
sd=[sd1 sd2];
var=sd.^2;
err0=sqrt(var./n);
ylabell=log(y);
for i=1:26
  err(i)=err0(i)*ylabell(i)/y(i);
end

%% Tracking model true data
u=utrue_smooth;
tplot=tdata;
ulabel=zeros(1,length(tlabel));
tlab=tlabel;
for i=1:length(tlabel)
  for t=1:length(tplot)
    if tplot(t)>min(tlab)
      ulabel(1,i)=u(1,t);
      tlab(1,i)=0;
      break
    end
  end
end
w=err.^-2;
```

```
W=diag(w);

X=[ones(length(tlabel),1) ulabel'-2.9559*ones(length(tlabel),1)];
bls=inv(X'*W*X)*X'*W*ylabell';
a1=bls(1);
b1=bls(2);
yhat1=a1+b1*(ulabel-2.9559*ones(1,26));

for t=1:length(u)
   yhatplot_true(t)=a1+b1*(u(t)-2.9559);
end

WMSE_Tracking=sum((ylabell-yhat1)*W*(ylabell-yhat1)')/sum(diag(W))

for j=1:N
  lnw(j)=log(sqrt(1/w(j)));
  z(j)=(ylabell(j)-yhat1(j)).^2*w(j);
end
AICc_Tracking_true=2*k+N*log(2*pi)+N+2*sum(lnw)+N*log(sum(z)/N)+2*k*(k+1)/(N-k-1)

%% Tracking model simulated data
k=7;
u=usim_smooth;
tplot=tdata;
ulabel_sim=zeros(1,length(tlabel));
tlab=tlabel;
for i=1:length(tlabel)
   for t=1:length(tplot)
      if tplot(t)>min(tlab)
         ulabel_sim(1,i)=u(1,t);
         tlab(1,i)=0;
         break
      end
   end
```

```
end
w=err.^-2;
W=diag(w);

X=[ones(length(tlabel),1) ulabel_sim'-2.9559*ones(length(tlabel),1) ];
bls_sim=inv(X'*W*X)*X'*W*ylabell';
a1_sim=bls_sim(1);
b1_sim=bls_sim(2);

yhat1_sim=a1_sim+b1_sim*(ulabel_sim-2.9559*ones(1,26));
for t=1:length(u)
   yhatplot_sim(t)=a1_sim+b1_sim*(u(t)-2.9559);
end

WMSE_Tracking_sim=sum((ylabell-yhat1_sim)*W*(ylabell-yhat1_sim)')/sum(diag(W))
for j=1:N
  lnw(j)=log(sqrt(1/w(j)));
  z(j)=(ylabell(j)-yhat1_sim(j)).^2*w(j);
end
AICc_Tracking_sim=2*k+N*log(2*pi)+N+2*sum(lnw)+N*log(sum(z)/N)+2*k*(k+1)/(N-k-1)

figure(4)
subplot(2,1,1)
yhat1=a1+b1*(ulabel-2.96*ones(1,26));
errorbar(ulabel,ylabell,err,'bo','LineWidth',1.0), hold on, grid
plot(ulabel,yhat1,'b','LineWidth',1.0), hold off
axis([2.95 3.03 2.12 2.33])
xlabel('d^1^8O_s_m_o_o_t_h')
ylabel('log mean dorsal fin ray number')
subplot(2,1,2)
yhat1=a1_sim+b1_sim*(ulabel_sim-2.96*ones(1,26));
errorbar(ulabel_sim,ylabell,err,'bo','LineWidth',1.0), hold on, grid
plot(ulabel_sim,yhat1_sim,'m','LineWidth',1.0), hold off
axis([2.95 3.03 2.12 2.33])
```

```
xlabel('d^1^8O_s_i_m_, _s_m_o_o_t_h')
ylabel('log mean dorsal fin ray number')

%% WLS model
k=3;
X=[ones(26,1) tlabel'];
bls=inv(X'*W*X)*X'*W*ylabell';
a2=bls(1);
b2=bls(2);
for i=1:26
   yhat2(i)=a2+b2*tlabel(i);
end

WMSE_WLS=sum((ylabell-yhat2)*W*(ylabell-yhat2)')/sum(diag(W))

for j=1:N
  lnw(j)=log(sqrt(1/w(j)));
  z(j)=(ylabell(j)-yhat2(j)).^2*w(j);
end
AICc_WLS=2*k+N*log(2*pi)+N+2*sum(lnw)+N*log(sum(z)/N)+2*k*(k+1)/(N-k-1)

figure(5)
errorbar(tlabel,ylabell,err,'bo','LineWidth',1.0), hold on
plot(tplot,yhatplot_true,'b','LineWidth',1.0)
plot(tlabel,yhat2,'--b','LineWidth',1.0)
plot(tplot,yhatplot_sim,'m','LineWidth',1.0)
hold off, grid
axis([-10 -9.88 2.12 2.33])
xlabel('Million years')
ylabel('log mean dorsal fin ray number')

figure(6)
plot(tdata,utrue_smooth,'LineWidth',1.0),grid
axis([-10.1 -9.8 2.88 3.08]);
```

```
ylabel('d^1^8O_s_m_o_o_t_h')
xlabel('Million years')

%% Akauke weights

 D_WLS=-121+132;
 D_true=-131+132;
 D_sim=-122+132;

 L_WLS=exp(-0.5*D_WLS);
 L_true=exp(-0.5*D_true);
 L_sim=exp(-0.5*D_sim);

 Aweight_WLS=L_WLS/(1+L_WLS+L_true+L_sim)
 Aweight_true=L_true/(1+L_WLS+L_true+L_sim)
 Aweight_sim=L_sim/(1+L_WLS+L_true+L_sim)
 Aweight_sinus=1/(1+L_WLS+L_true+L_sim)

 SUM_weights=Aweight_WLS+Aweight_true+Aweight_sim+Aweight_sinus
```
